# Global mistranslation facilitates sampling of beneficial mutations under stress

**DOI:** 10.1101/650846

**Authors:** Laasya Samhita, Parth K Raval, Deepa Agashe

**Author notes:** Correspondence: Laasya Samhita (,), Deepa Agashe.

## Abstract

Mistranslation is typically deleterious, but can sometimes be beneficial. Although a specific mistranslated protein can confer a short-term benefit in a particular environment, the prevalence of high global mistranslation rates remains puzzling given the large overall cost. Here, we show that generalized mistranslation enhances early *E. coli* survival under various forms of DNA damage, because it leads to early activation of the DNA damage-induced SOS response. Mistranslating cells therefore maintain larger populations, facilitating later sampling of critical beneficial mutations. Thus, under DNA damage, both basal and induced mistranslation (through genetic or environmental means) increase the number of genetically resistant and phenotypically persistent cells. Surprisingly, mistranslation also increases survival at high temperature. This wide-ranging stress resistance relies on Lon protease, which is revealed as a key effector that induces the SOS response in addition to alleviating proteotoxic stress. The new links between error-prone protein synthesis, DNA damage, and generalised stress resistance indicate surprising coordination between intracellular stress responses, and suggest a novel hypothesis to explain high global mistranslation rates.

## INTRODUCTION

The rate of protein mistranslation is amongst the highest known error rates in cellular biosynthetic processes, ranging from 1 in 10,000 to 1 in 100 mis-incorporated amino acids in *E.coli* ^1,2^. As a result, 10 to 15% of all proteins in an actively growing *E.coli* cell are likely to carry at least one mis-incorporated amino acid ^3,4^, implying a high tolerance for mistakes. This is puzzling because mistranslation is thought to be deleterious, and cells have evolved several proofreading mechanisms to minimise error reviewed in ^5^. Counterintuitively, a body of work showing that cells elevate basal mistranslation levels under specific stresses reviewed in ^6,7^ suggests that high mistranslation may also evolve under positive selection. This is further supported by multiple examples of the selective advantage of specific mistranslated proteins. For instance, in *Mycobacterium smegmatis*, increasing specific amino acid substitutions at glutamate and aspartate tRNAs generates a mixed population of wild type and mistranslated RNA polymerase molecules ^8^. The resulting amino acid substitutions inhibit RNA polymerase activity and increase resistance to rifampicin (an antibiotic that targets RNA polymerase). However, it remains unknown whether selection favouring specific mistranslated proteins in distinct environments is sufficient to drive increased global mistranslation rates.

Alternatively, selection may directly favour high global mistranslation rates by generating a “statistical proteome” – a bet-hedging strategy where a few cells with specific mistranslated proteins can survive a given environmental stress ^9,10^. The only natural (non-manipulated) example of such general proteome-wide beneficial mistranslation comes from fascinating work on mis-methionylation in *E.coli*. In anaerobic environments or upon exposure to low concentrations of chloramphenicol, the methionyl tRNA synthetase enzyme loses its succinyl modifications, reducing enzyme fidelity ^11^. As a result, the enzyme amino-acylates methionine onto non-cognate tRNAs, causing ‘mis-methionylation’ ^12^ and increasing survival under anaerobic and antibiotic stress. However, we do not yet know why the succinyl modifications are altered under these specific stresses, nor the underlying mechanism. More generally, increasing overall mistranslation levels is typically deleterious, reviewed in ^7,13^, suggesting a narrow range of error rates in which the potential benefit of a few specific mistranslated proteins could outweigh the larger overall cost of mistranslated proteins.

Here, we propose a new hypothesis that bypasses the need for specific mistranslated proteins, making a broad fitness benefit of global mistranslation plausible. We demonstrate a mechanism by which generalized mistranslation increases resistance to multiple stresses in *E. coli*. To mimic natural cellular responses to environmental stress, we initiated our study using a strain with genetically depleted initiator tRNA (tRNAi) content (henceforth “Mutant”, carrying only one of four wild type “WT” tRNAi genes ^14^). As central players in translation, cellular tRNA levels have a major impact on mistranslation ^15,16^, and are rapidly altered in response to environmental change ^17–19^. Initiator tRNA (tRNAi) levels are especially interesting because translation initiation is a rate limiting step ^20^, and tRNAi levels change under various stresses. For instance, in *E.coli*, amino acid starvation is accompanied by a transcriptional tRNAi downregulation during the stringent response ^21^, while mammalian cells reduce tRNAi levels on exposure to stressors such as the toxin VapC ^22^ and high temperature ^23^. Depletion of tRNAi causes at least one kind of mistranslation, allowing promiscuous non-AUG initiation by elongator tRNAs ^16,22^. We therefore tested whether mistranslation resulting from tRNAi depletion in the Mutant leads to a general survival advantage.

We first carried out a Biolog screen ^24^ comparing WT and Mutant growth across a range of environments, including 48 antibiotics with various modes of action. The Mutant showed higher growth in the presence of Novobiocin (Fig. S1), a fluoroquinolone antibiotic that inhibits DNA gyrase and causes DNA damage. Further work showed that inducing mistranslation via multiple mechanisms conferred protection against several kinds of DNA damage, via induction of the well-studied bacterial SOS response. Increased mistranslation brings cells closer to the intracellular molecular threshold for SOS induction, such that mistranslating cells sense and repair DNA damage sooner than the wild type. The resulting increase in early survival facilitates the eventual emergence of genetic resistance as well as phenotypic persistence under antibiotic stress. Interestingly, the mistranslation-induced SOS response is also beneficial in other conditions, increasing persistence and survival at elevated temperature. Thus, we have uncovered a general, novel link between mistranslation and DNA damage that integrates two major cellular pathways and suggests a new hypothesis for the evolution of mistranslation rates.

## RESULTS

### Mistranslation increases resistance to DNA damage by enhancing early cell survival

Compared to WT, the mistranslating Mutant with depleted tRNAi showed higher survival under DNA damage of various kinds, induced by exposure to UV radiation (base dimerization), hydrogen peroxide (base oxidation) or the antibiotic ciprofloxacin (‘Cip’, a more potent DNA gyrase inhibitor than Novobiocin, that causes double stranded DNA breaks) (Fig. 1a–c). Higher Cip resistance in the Mutant did not arise as a by-product of slower growth (in LB, the Mutant has a doubling time of ~1.0 h compared to ~0.6 h for WT; Fig. S2a): WT Cip resistance did not increase when grown in glycerol, where it has a 5-fold lower doubling time (Fig. S2b). These results were intriguing because mistranslation has no known connection with DNA damage or its repair. To determine the mechanisms underlying this connection, we focused on Cip resistance.

**Figure 1.**
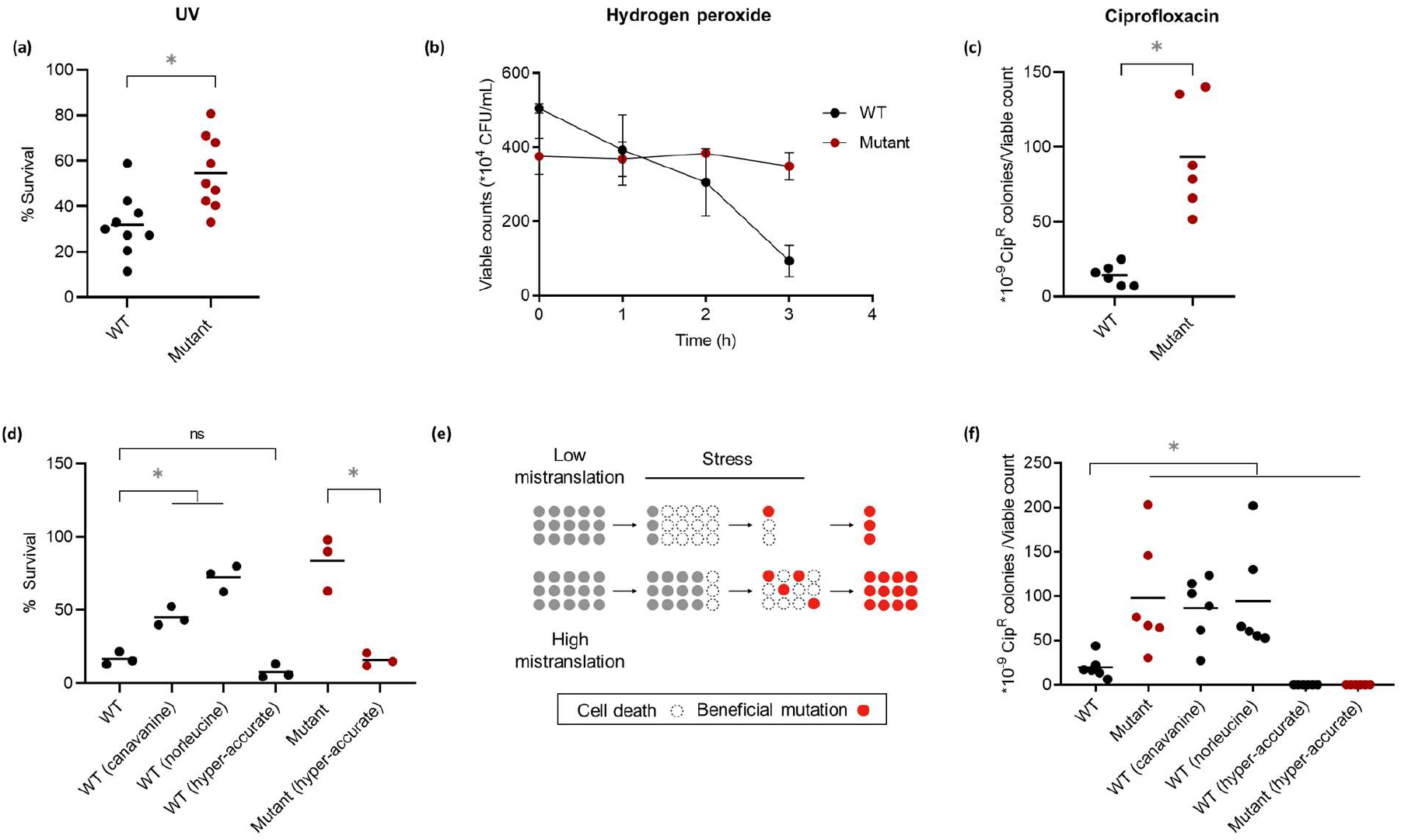
Mistranslation confers resistance to DNA damage by increasing early survival. (a) Survival of WT and Mutant mid log phase cultures (OD_600nm_ ~0.6, n=9) exposed to 20 J/m^2^ of UV-C radiation for 5 s. The plot shows mean % survival relative to the number of colonies on a non-irradiated control plate. Mann-Whitney U test, Mutant>WT, U=9.5, P=0.0046 (b) Time course of survival (viable counts) of mid log phase cultures of WT and Mutant (n=4) treated with 5 mM hydrogen peroxide. At 3 h, Mutant>WT, Mann-Whitney U test, U=0, P=0.029 (c) Resistance of WT and Mutant mid log phase cultures inoculated from single colonies (n=6), pulsed with 20 ng/mL ciprofloxacin (Cip) for 1 h, and plated on LB agar plates with vs. without 50 ng/mL Cip (Cip50). The plot shows the average proportion of resistant colonies relative to total viable counts. Mann-Whitney U test, U=0, P=0.002 (d) Early survival (tolerance) of WT, mistranslating and hyper-accurate strains treated with Cip50 for 2 h, from mid log phase cultures inoculated from single colonies (n=3) treated with Cip50 for 2 h. We estimated viable counts before and after exposure. The plot shows mean % survival in each case. t tests: Mutant>WT, t=6.13, P=0.02; WT(canavanine)>WT, t=6.1, P=0.005; WT(norleucine)>WT, t=9.6, P=0.002; WT(hyper-accurate) vs. WT, ns, t=2.3, P=0.08; Mutant(hyper-accurate)<Mutant, t=6.2, P=0.02 (e) Schematic of the proposed model of mistranslation leading to increased sampling of beneficial mutations via enhancement of early survival. (f) Resistance of WT, mistranslating and hyper-accurate strains to Cip50, from mid log phase cultures inoculated from single colonies (n=6), pulsed with Cip20 for 1 h, and plated on Cip50 LB agar. The plot shows the mean proportion of resistant colonies relative to total viable counts. Mann-Whitney U tests: Mutant>WT: U=1, P=0.0043; WT(canavanine)>WT: U=1, P=0.0043; WT(norleucine)>WT: U=0, P=0.0022; WT(hyper-accurate)<WT: U=0, P=0.0022; Mutant(hyper-accurate)<Mutant: U=0, P=0.0022; Mutant(hyper-accurate)<WT: U=0, P=0.0022. Asterisks indicate a significant difference.

Whole genome sequencing showed that each Cip-resistant (Cip^R^) colony of WT and Mutant (after 24 h on Cip plates) had a single mutation within the well-known QRDR (Quinolone Resistance Determining Region) of the *gyrA* gene (Table S1). Thus, while WT and Mutant cells acquired identical beneficial mutations, the Mutant was more likely to sample them. However, WT and Mutant had similar basal mutation frequency (Fig. S3), suggesting that higher mutation rate could not explain higher Cip resistance in the Mutant. Instead, we found that the Mutant had greater early survival after Cip exposure (after 2 h; Fig.1d), and cells sampled at this point did not have any QRDR mutations. Therefore, this early survival was not due to genetic resistance, but implies a form of tolerance. The ~5 fold difference in population size meant that a higher proportion of cells in Mutant cultures could sample *gyrA* mutations, ultimately increasing Cip resistance. Together, these results suggested that mistranslation indirectly enhanced Cip resistance by increasing early survival (Fig. 1e).

To test the generality of this result, we manipulated mistranslation levels by (a) reducing global mistranslation via hyper-accurate ribosomes (Methods; Fig. S4) and (b) increasing WT mistranslation by adding the non-proteinogenic amino acids canavanine or norleucine to the growth medium ^25,26^. Increasing mistranslation rates consistently increased early survival (Fig. 1d) and Cip resistance in WT (Fig. 1f), whereas suppressing basal mistranslation decreased early survival and Cip resistance in both WT and Mutant (Fig. 1d and 1f). We next focused on understanding the mechanistic basis of this effect.

### Mistranslation mediates ciprofloxacin resistance via the SOS response

In response to DNA damage, bacterial cells induce the SOS response, which controls the expression of several DNA repair pathways ^27^. Briefly, DNA damage generates single stranded DNA that binds to the protein RecA. Activated RecA stimulates cleavage of LexA (a repressor), which in turn induces the SOS response, de-repressing several DNA repair genes (Fig. 2a). When we blocked SOS induction by replacing the WT *lexA* allele with a non-degradable allele lexA3; ^28^ and challenged cultures with Cip, both WT and Mutant showed low early survival (Fig. S5) and negligible Cip resistance (Fig. 2b), as expected in the absence of an intact DNA repair response. Thus, mistranslation-induced increase in tolerance leading to Cip resistance depends on the SOS response.

**Figure 2.**
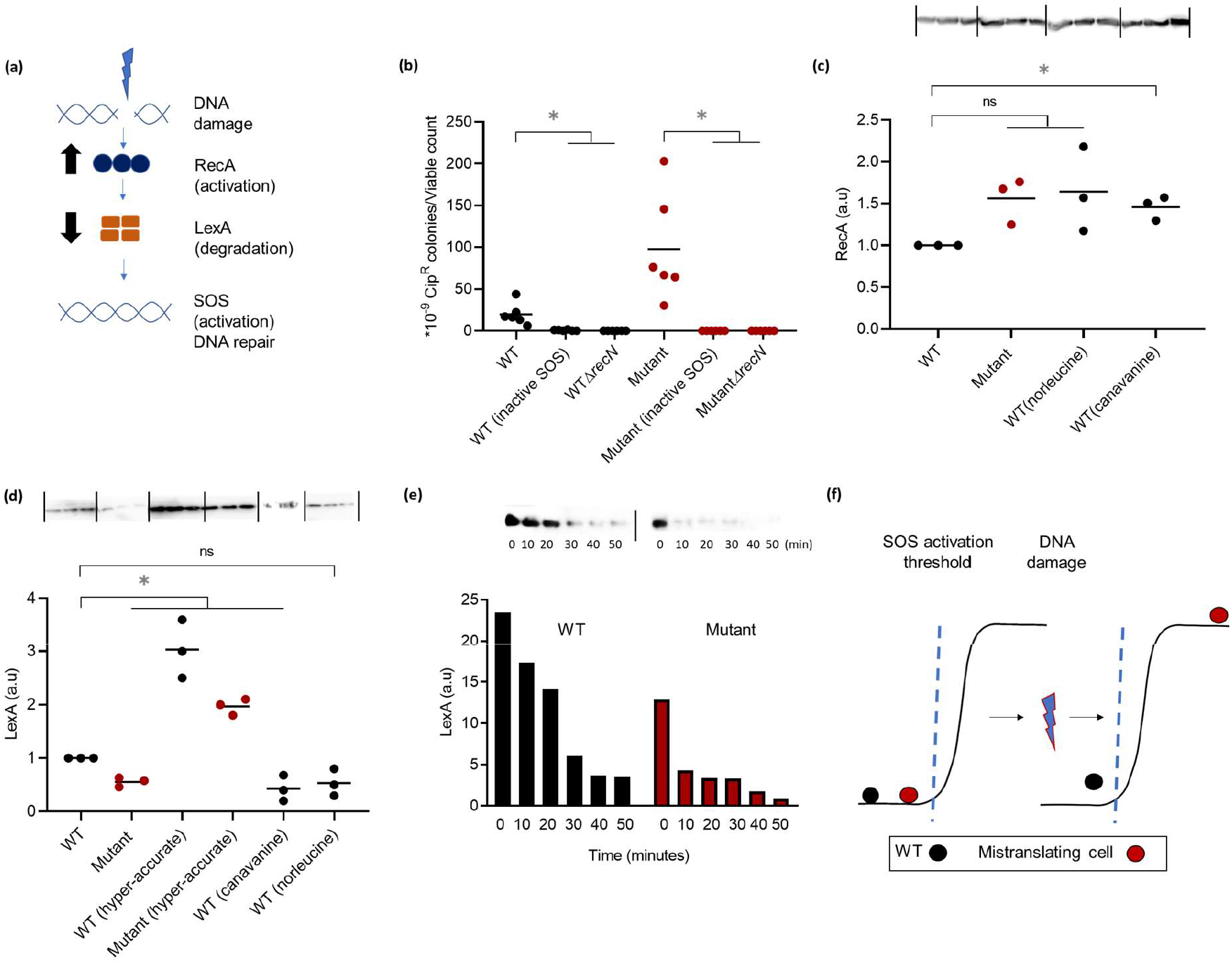
Mistranslation mediates ciprofloxacin resistance via the SOS response. (a) Schematic of the SOS response in *E.coli* (b) Survival of WT and Mutant mid-log phase cultures (OD_600nm_~0.6) from single colonies (n=6) pulsed with 20 ng/mL ciprofloxacin (Cip20) for 1 h and plated on LB agar with vs. without 50 ng/mL Cip (Cip50). The plot shows the mean proportion of resistant colonies relative to total viable counts. SOS was inactivated using the *lexA3* allele. Mann-Whitney U test: Mutant>WT, U=1, P=0.0043; WT(inactive SOS)<WT, U=0, P=0.0022; Mutant(inactive SOS)<Mutant, U=0, P=0.0022; WTΔ*recN*<WT, U=0, P=0.0022; MutantΔ*recN*<Mutant, U=0, P=0.0022 (c) A representative blot showing RecA protein levels from mid log phase cultures of WT and mistranslating strains (n=3). Quantification of mean blot band/total protein is represented in arbitrary units, relative to WT. Paired t tests: WT vs. Mutant, ns, t=3.5, P=0.07; WT(canavanine) >WT, t=5.1, P=0.03; WT vs WT(norleucine), ns, t=2.2, P=0.1 (d) A representative blot showing LexA protein levels from mid log phase cultures of WT and mistranslating strains (n=3). Quantification of mean blot band/total protein is represented in arbitrary units, relative to WT. Paired t tests: Mutant<WT, t=8.7, P=0.01; WT(canavanine)<WT, t=29.6, P<0.0001; WT vs WT(norleucine), ns, t=3.4, P=0.07; WT(hyper-accurate)>WT, t=12.4, P=0.006; Mutant(hyper-accurate)> Mutant, t=10.4, P=0.009 (e) Time course of LexA degradation from 0 to 50 min post exposure to Cip20. One blot and bands normalised to total protein are shown here; for more experimental blocks, see Fig. S6 (f) Schematic of the proposed model of the state of WT and mistranslating strains with respect to the SOS activation threshold. Asterisks indicate a significant difference between strains.

The SOS response has two opposing aspects: rapid DNA repair, and increased mutagenesis due to the activation of error-prone polymerases. The latter temporarily elevates mutation rate, increasing the supply of beneficial mutations ^29^. However, as mentioned above, WT and Mutant had similar basal mutation frequencies (Fig. S3). Therefore, we reasoned that the increased early survival of mistranslating strains must be aided by faster or more efficient repair and recombination. To test this, we deleted RecN – a key member of the SOS-linked recombination mediated repair pathway ^30^. The deletion led to decreased early survival upon Cip exposure (Fig. S6) and a complete loss of Cip resistance (Fig. 2b), indicating that repair and recombination functions indeed underlie the increased Cip resistance observed in the mistranslating Mutant.

### Mistranslation enhances Cip resistance by allowing faster induction of the SOS response

Since both WT and Mutant rely on the SOS response for Cip resistance, the ~5-fold greater survival of the Mutant (Fig. 1d) continued to be a puzzle. We hypothesized that the survival advantage arose from differential induction of SOS due to mistranslation, allowing rapid DNA repair. Consistent with this hypothesis, greater mistranslation was associated with slightly (though not significantly) higher basal RecA levels across multiple experimental blocks (Fig. 2c). These results suggest that even in the absence of DNA damage, RecA was already elevated in mistranslating strains, positioning the cell closer to the SOS induction threshold (Fig. 2a and 2f). To test this, we induced the SOS response in WT and Mutant and monitored the time course of LexA degradation. The Mutant degraded LexA within 10–20 minutes of SOS induction, while the WT took an additional 10–20 minutes (Fig. 2e and Fig. S7). Similarly, LexA was degraded at lower concentrations of Cip in the Mutant (Fig. S8). Note that the Mutant is not already ‘stressed’ and has similar basal LexA levels to the WT (Fig. S8); it is only upon encountering DNA damage that LexA starts degrading.

Together, these observations suggest that in mistranslating strains, (i) LexA is degraded faster upon encountering the DNA damaging stress, and (ii) LexA is degraded at a lower magnitude of the stress (Fig. 2f). Thus, we demonstrate a direct causal relationship between mistranslation, induction of the SOS response, and enhanced survival under Cip.

### Mistranslation induces the SOS response via Lon protease

In our experiments, we induced mistranslation in distinct ways, and consistently observed increased survival under DNA damage. Canavanine and norleucine respectively replace arginine and leucine in the proteome, whereas tRNAi depletion causes mis-initiation with elongator tRNAs. The parallel outcomes from these diverse modes of mistranslation suggested a general mechanistic link between mistranslation and SOS response. Based on prior studies, we suspected that Lon – a key protease across eubacteria – may represent such a link. In mistranslating *E.coli* cells, Lon alleviates the associated deleterious effects by degrading aggregated and non-functional proteins ^3^. In *Pseudomonas*, Lon is essential for RecA accumulation and induction of the SOS response, and is suggested to degrade RecA repressors such as RecX and RdgC ^31^.

We therefore hypothesized that our mistranslating strains may have higher amounts of Lon, in turn accumulating RecA and bringing cells physiologically closer to the threshold for SOS induction. Because Lon is part of the *E. coli* heat shock regulon ^32^, we also suspected a general increase in the heat shock response. Indeed, mistranslating cells had higher levels of Lon protease (Fig. 3a), as well as the heat shock transcription factor sigma 32 (Fig. S9). For technical reasons, we were unable to knock out Lon in our wild type strain KL16. Hence, we deleted Lon in *E. coli* MG1655. While MG1655 had comparable ciprofloxacin resistance to our WT (KL16; Fig. S10), deleting Lon decreased Cip resistance (Fig. 3b) and increased LexA levels in SOS-induced cells (Fig. 3c). Conversely, over-expressing Lon enhanced early survival (Fig. S11) and resistance to Cip (Fig. 3b), and reduced LexA levels in both WT and Mutant upon SOS induction (Fig. 3c). Over-expressing Lon also elevated basal RecA levels (in the absence of any DNA damage), further supporting our hypothesis (Fig. S12). Finally, since Lon is part of the heat shock regulon, we predicted that prior exposure to high temperature should induce Lon and increase resistance to DNA damage. True to expectation, cells grown at 42°C for three hours had higher Cip resistance (Fig. 4a). Together, these results strongly support a key role for Lon in mediating Cip resistance by inducing the SOS response.

**Figure 3.**
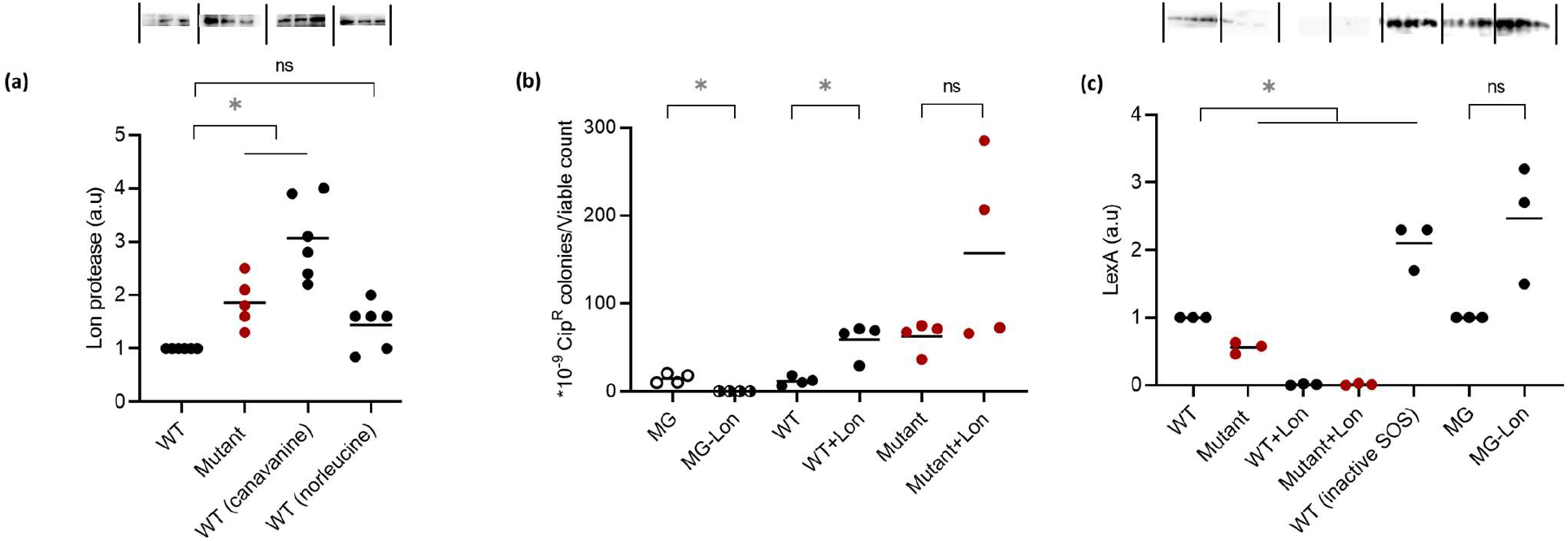
Mistranslation induces the SOS response via Lon protease. (a) A representative blot showing Lon protein levels from mid log phase cultures (OD_600nm_ ~0.6) of WT and mistranslating strains (n=3). Quantification of mean blot band/total protein is represented in arbitrary units, relative to WT. Paired t tests: Mutant>WT, t=4.2, P=0.01; WT(canavanine)>WT, t=6.5, P=0.003; WT vs. WT(norleucine), ns, t=2.5, P=0.05 (b) Survival of MG1655, MGΔ*lon* (MG-Lon), WT (KL16), WT+Lon, Mutant and Mutant+Lon on Cip50, from mid-log phase cultures from single colonies (n=4) pulsed with Cip20 for 1 hr and plated on LB agar with vs. without Cip50. The plot shows the mean proportion of resistant colonies relative to total viable counts. Paired t tests: MGΔ*lon*<MG, t=5.3, P=0.01; WT+Lon>WT, t=5.5, P=0.01; Mutant vs. Mutant+Lon, ns, t= 1.9, P=0.14 (c) A representative blot showing LexA protein levels from mid log phase cultures of WT and mistranslating strains (n=3). Quantification of mean blot band/total protein is represented in arbitrary units relative to WT. SOS was inactivated using the *lexA3* allele. Paired t tests: Mutant<WT, t=8.8, P=0.01; WT+Lon<WT, t=171.5 P<0.0001; Mutant+Lon<Mutant, t=10.4, P=0.009; WT(inactive SOS)>WT, t=5.5, P=0.03; MG vs MGΔ*lon*, ns, t=3.2, P=0.08. Asterisks indicate a significant difference between strains.

**Figure 4.**
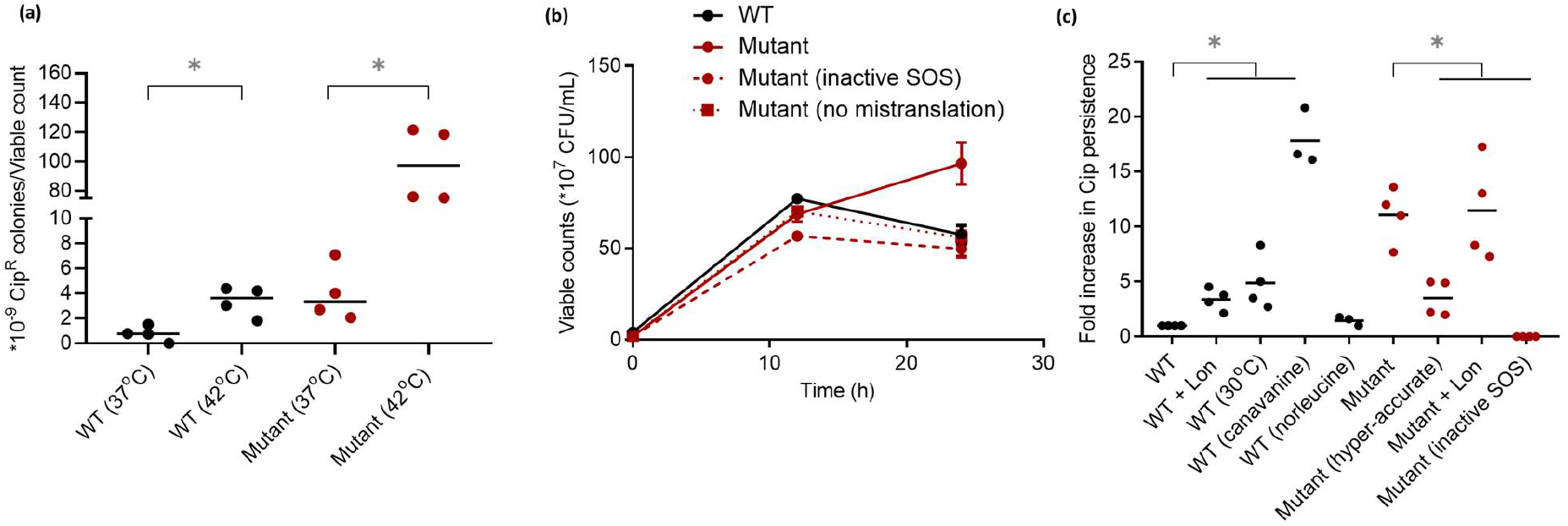
Mistranslation-induced SOS response enhances survival in other stresses. (a) Survival of WT and Mutant mid log phase cultures (OD_600nm_~0.6) grown overnight at 37°C from single colonies (n=4), sub-cultured and grown at 37°C or 42°C (to induce the heat shock response while cells entered log phase) for 3.5 hours, and plated on LB agar with vs. without 50 ng/mL Cip (Cip50). The plot shows the mean proportion of resistant colonies relative to total viable counts. Mann-Whitney tests: WT(42°C)>WT(37°C), U=0, P=0.0286; Mutant(42°C)>Mutant(37°C), U=0, P=0.0286 (b) Total viable counts of various strains at 0, 12 and 24 h after exposure to 42° C. SOS was inactivated using the *lexA3* allele. At 24h, Mann-Whitney tests: Mutant>WT, U=0, P=0.0286; Mutant(inactive SOS) vs. WT, ns, U=4.5, P=0.4; Mutant(hyper-accurate) vs. WT, ns, U=8, P>0.99 (c) LexA levels from mid log phase cultures (n=3) grown at 42°C for 12 h. SOS was inactivated using the *lexA3* allele. Quantification of mean blot band/total protein is shown in arbitrary units relative to WT. Paired t test, Mutant<WT, t=5.3, P=0.03. Asterisks indicate a significant difference between strains.

### Mistranslation-induced SOS response enhances survival in other stresses

As mentioned above, Lon is part of the heat shock regulon; hence we tested whether mistranslation also increased survival under high temperature. As predicted, we found that the Mutant had greater survival at high temperature, especially in the stationary phase of growth (after 12 hours; Fig 4b). We also observed this growth advantage in WT cells treated with norleucine and canavanine, although canavanine results were variable (Fig. S13). Importantly, Lon alone could not explain the greater survival at high temperature: it required both mistranslation and a functional SOS response (Fig. 5b); Lon levels at 42°C were comparable across WT and Mutant (Fig. S14); and LexA was degraded in the Mutant but not in the WT (Fig. S15). Interestingly, RecN is important for high temperature survival of both WT and Mutant, suggesting that survival is influenced by functional DNA repair (Fig. S16). The clear dependence of the survival advantage on SOS induction suggests cross-talk between mistranslation, the heat shock response and SOS induction.

**Figure 5.**
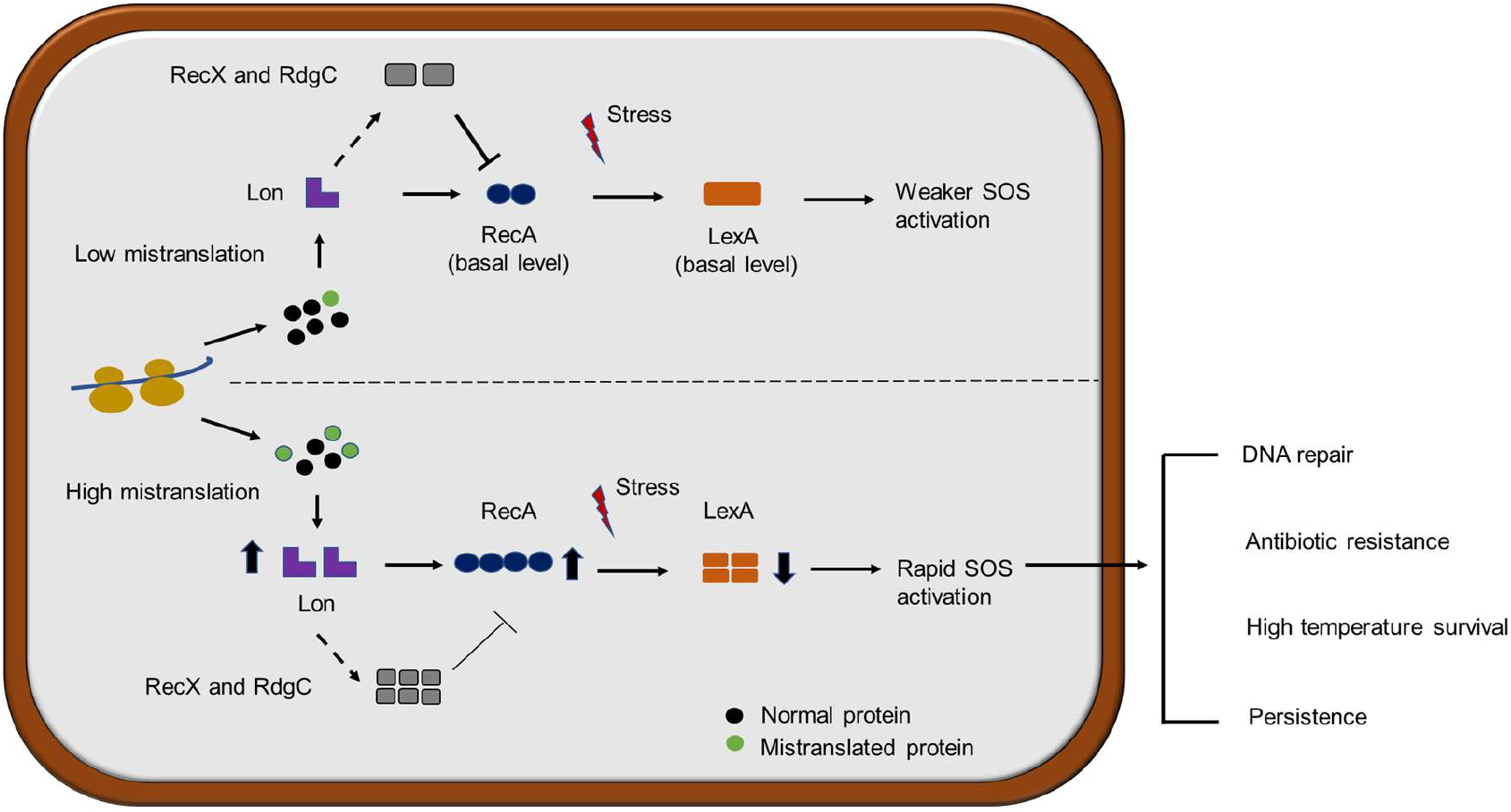
Proposed model for stress resistance mediated by faster SOS activation in mistranslating strains.

The SOS response controls over 40 genes in *E.coli*, including five toxin-antitoxin modules. Of these, the TisA/TisB module induces persistence (a state of metabolic dormancy leading to increased tolerance to antibiotics and other drugs) by reducing the proton motive force across the cell membrane ^33^. We therefore tested whether inducing mistranslation increases persistence following exposure to Cip and other antibiotics. We defined the number of persisters as the number of colony forming units that survive exposure to a lethal antibiotic concentration (Cip 200), but do not divide actively in its presence. We found that mistranslation increased persistence to Cip (Fig. 4c). In yeast, DNA damage also induces persisters that carry more mutations ^34^, as expected under an elevated SOS response leading to mutagenesis. However, in our case – true to the conventional understanding of persistence – we did not find any mutations when we sequenced whole genomes of WT or Mutant persister cells, as well as their hyper-accurate counterparts.

Our results thus show that mistranslation-induced early activation of the SOS response is responsible for multiple stress resistance phenotypes. We suggest a model whereby mistranslation of various kinds increases Lon protease levels, triggering an early induction of the SOS response that enhances cell survival not only under DNA damage, but also under other stresses (Fig. 5).

## DISCUSSION

High global translation error rates have remained an enduring puzzle, given the large overall costs of generalized mistranslation compared to the benefits of generating specific mistranslated proteins in particular environments. In particular, this problem has mired the hypothesis that mistranslation rates may evolve under positive selection. Here, we diminish this barrier by demonstrating that basal as well as induced non-specific mistranslation enhances early survival under DNA damage by rapidly inducing the SOS response, making a larger pool of cells available for subsequent genetic change. We further show that this effect is mediated by increased levels of a key protease that deals with aberrant proteins as well as several important regulatory enzymes. Since survival does not rely on the chance generation of specific mistranslated proteins, cells effectively bypass the deleterious load of high mistranslation, and instead use it to trigger stress responses and alleviate damage to cellular components. Most stresses that are commonly encountered by bacteria (oxidative stress, high temperature, radiation, starvation) induce mistranslation (through damage to proteins), DNA damage, or both. Hence, it seems fitting that these two phenomena should be linked, and should operate through a common effector molecule.

Although prior work had suggested a link between mistranslation and SOS response in ageing bacterial colonies ^35^, the underlying mechanism and generality of the proposed link remained unknown. In contrast to the known mutagenic impact of SOS induction (generating “hopeful monsters”), we demonstrate that mistranslation is generally beneficial under stress because it enhances the rapid repair modules controlled by the SOS response. Our proposed model (Fig. 5) thus lends support to the hypothesis that global mistranslation levels could evolve under positive selection. More broadly, our results imply that generalized non-genetic changes can facilitate subsequent genetic adaptation by increasing short-term survival. This has been an attractive hypothesis with limited and protein-specific prior support. In *S. cerevisiae*, ribosomal frameshifting alters localization of a specific protein, a phenotype that is then fixed by genetic mutations in a few generations ^36^. Similarly, changes in Hsp90 levels in *Candida albicans* ^37^ and at least one phenotype conferred by the prion PSI+ in wild yeasts can be stabilised over evolutionary time ^38^. Our results generalize these effects, providing clear evidence that non-directed phenotypic changes can facilitate improved stress resistance by enhancing the subsequent sampling of beneficial mutations.

Our work also helps to synthesize a diverse body of prior results into a cohesive framework. It is well known that mistranslation may lead to misfolding and protein aggregation, which is typically deleterious reviewed in ^39^. However, a recent study showed that cells carrying intracellular protein aggregates were more stress-resistant ^40^. Interestingly, these cells had upregulated chaperones such as DnaK and proteases such as ClpP, suggesting that protein aggregation can also precipitate stress resistance. Prior work also shows that Lon is required to alleviate the toxic effects of mistranslation ^3^; but except for one study in *S. cerevisiae* ^41^, there was no evidence linking the heat shock response with mistranslation. We show that the heat shock response is activated by general mistranslation (Fig. S9), leading to increased Lon levels. Our model could also explain the puzzling observation that Lon protease function determines sensitivity to high concentrations of quinolone antibiotics (which target DNA gyrase, with no direct connection to Lon) such as nalidixic acid ^42^ and levofloxacin ^43^. Our results show that impairing Lon hinders cells’ ability to rapidly induce the SOS response and repair damaged DNA. The central role of Lon in bridging mistranslation and the SOS response is also supported by previously observed links between the heat shock response and the SOS response. For example, in *Listeria monocytogenes*, heat shock directly triggers the SOS response ^44^. In *E. coli*, the heat shock chaperone GroE also induces expression of the mutator polymerase UmuD, hitherto thought to be regulated only via the SOS response ^45^. Finally, when exposed to levofloxacin, *E.coli* cells express both the SOS response and heat shock genes ^43^. These results corroborate our observation that mistranslation and SOS activation together increase heat resistance, though we do not yet know precisely how this phenotype is regulated. Altogether, we suggest that Lon acts as a key molecule that coordinates several aspects of stress responses, encompassing toxin-antitoxin regulation, survival under anaerobic conditions, SOS, heat shock, antibiotic resistance, and cell division reviewed in ^32^.

The SOS response is among the best studied pathways in *E.coli*, inducing DNA repair genes in response to double strand breaks and stalled replication forks generated by severe DNA damage. Yet, we continue to unravel new phenotypes controlled by this response. Instead of being directed solely at DNA repair, the SOS response is turning out to be central for several stressful situations. Similarly, the causes and impacts of mistranslation also continue to be extensively explored, with new details surfacing each day. At the moment, we cannot determine whether mistranslation was co-opted by the DNA damage response as a trigger, or vice-versa. Irrespective of which response evolved first, it is clear that diverse cellular mechanisms are linked in unexpected ways, co-ordinating the cellular response to multiple stresses. While these phenomena are independently well studied, we can now connect them using a single effector molecule, Lon protease. Our study raises the question of whether such novel links could themselves be evolving in different directions, leading to cross talk between mutation-independent phenotypic variation and genetic change in response to stress.

## ACKNOWLEDGEMENTS

We thank Anjana Badrinarayanan, Sunil Laxman, and members of the Agashe lab for discussion and critical comments on the manuscript. We thank Awadhesh Pandit and Tejali Naik for help with whole genome sequencing at the NCBS sequencing facility; Dipankar Chatterjee and Kuldeep Gupta (Indian Institute of Science, Bangalore) for access to and help with their Biolog machine; Kurt Fredrick (Ohio State University) for the luciferase assay system; and Jayaraman Gowrishankar and Nalini Raghunathan (Centre for DNA Fingerprinting and Diagnostics, Hyderabad) for the *lexA3* allele. We acknowledge funding and support from the Wellcome Trust-DBT India Alliance (grant IA/E/14/1/501771 to LS and grant IA/I/17/1/503091 to DA); the Indian Council for Medical Research (ICMR fellowship 3/1/3/JRF-2015 to PR); the Council for Scientific and Industrial research (CSIR research grant 37(1629)/14/EMR-II to DA); and the National Centre for Biological Sciences (NCBS-TIFR).

## AUTHOR CONTRIBUTIONS

LS and DA conceived the project; LS and DA designed experiments; LS and PR conducted experiments; LS and DA analysed data; LS and DA wrote the manuscript.

