## Supplementary material for "Global mistranslation facilitates sampling of beneficial mutations under stress"

**SUPPLEMENTARY INFORMATION**

**MATERIALS AND METHODS**

**Strains and growth media**

We used *E. coli* KL16 (“wild type”, WT) ^1^ and its derivatives for all experiments, and introduced mutations by P1 transduction (see below). The Mutant strain KL16*ΔZWV* (“Mutant”) lacks three of the four initiator tRNA (tRNAi) genes in *E.coli* and shows non-AUG initiation ^2^. We used P1 lysate raised on the KL16*ΔZWV::Kan* strain (a gift from Prof. Umesh Varshney at the Indian Institute of Science, Bangalore) to knock out the three tRNAi genes *metZ, metW* and *metV* (which are organised in an operon; ^3^) using P1 transduction. We removed the kanamycin marker inserted during this process using the plasmid pcp20 ^4^, so that our results were not confounded by the presence of an antibiotic cassette. We obtained the *lexA3* allele ^5^ that abolishes the SOS response from Dr Gowri Shankar at CDFD, Hyderabad; the S12 K43R mutation from the strain SS3242 from CGSC, Yale university; and the CGSC *recN* knockout strain JW541-1 (Keio collection) from Dr Anjana Badrinarayanan at NCBS, Bangalore. We used P1 transduction to introduce the relevant mutation (or gene knockout) into our WT or Mutant strains. Strain SS3242 contains a mutation in the protein S12 (K42R) that increases translation accuracy by reducing the frequency of decoding errors ^6^. Hence, we used this mutation to generate “hyper-accurate” strains with reduced overall mistranslation rates. We observed an ~10-fold increase in translation accuracy both in the WT and in the Mutant, quantified using a plasmid-based assay system that measures missense suppression in a firefly luciferase gene (gifted by Prof Kurt Fredrick, Ohio State University) (Fig. S4 and SI methods). To manipulate expression of Lon protease, we cloned the *lon* gene into the Tet^R^ low copy plasmid pACDH ^7^ using the restriction enzymes NotI and EcoRI (restriction sites italicized) and the following primer sequences: fp 5’ CTCAT*GCGGCCGC*GGAAATTAAACTAAGAGAGAGCTCT 3’ and rp 5’ CTCAT*GAATTC*TGCCAGCCCTGTTTTTATTAGT 3’and transformed WT and Mutant (carrying intact genomic copies of *lon*) with the plasmid. We grew bacterial cultures in LB or LB-agar plates containing 1.8% agar (Difco). Where applicable, we used M9 minimal medium: 1X M9 salts prepared from 5X M9 stock powder (Difco), 0.5% glucose, 0.4% casamino acids (Difco, where specified), 1mM Cacl_2_, 2.5 mM MgSO_4_ and 0.01% Vitamin B1 (thiamine). When required, we supplemented the medium with Ciprofloxacin (Cip) at concentrations 10, 20, 50,100 or 200 ng/mL, Kanamycin (25 μg/mL), Tetracycline (7.5 μg /mL), Canavanine (3 mg/mL), Norleucine (2.25 μg/mL) or IPTG (1 mM). All antibiotics and chemicals were obtained from Sigma-Aldrich.

**Initial screening of WT and mutant strains using phenotype micro arrays**

To screen for phenotypic differences between the WT and the mutant across a range of environments, we employed phenotype microarrays ^8^. We grew the WT and mutant overnight in Difco LB medium. In accordance with the established protocol, we made cell suspensions with a transmittance of 81% in the IF0a medium supplied by Biolog, and added the redox indicator tetrazolium violet to a final concentration of 0.01%. We then inoculated 100 µL of the suspension into each well of the phenotype microarray 96 well plates. As bacteria respire, tetrazolium violet is reduced to a purple color, the intensity of which indicates bacterial growth. We incubated plates at 37°C for 48 h in a Biolog OmniLog incubator, and measured the dye intensity every 15 min, plotted as the area under the curve (AUC) in the Biolog parametric software. If the AUC of the test strain is more than that of the reference strain, it indicates that the test strain has enhanced respiration (readout for growth) compared with the reference strain. Since this is a qualitative estimate and baselines vary across experiments ^9^, we used the phenotype microarray to identify environments where WT and mutant may have differential survival and growth. We conducted all further experiments under standard laboratory conditions using single colonies and test tubes. We are grateful to Dr Dipankar Chatterjee and Dr Kuldeep Gupta from the Indian Institute of Science, Bangalore, for making their Biolog reader available and for help with setting up the experiments.

**P1 transduction**

We used P1 transduction to move specific alleles across genetic backgrounds. We pelleted cells from an overnight recipient culture and re-suspended in an equal volume of MC buffer (100 mM MgSO_4_.7H_2_O and 5 mM Cacl_2_.2H_2_O). We mixed 100 μl of cells with 50 μl of ~10^7^ pfu/mL phage lysate (made up to 100 μl with MC buffer), or with equal volumes of different dilutions of P1 lysate made in MC buffer (undiluted, 1:2 and 1:10). We incubated this mixture at 37^o^C for 20 minutes, pelleted cells by centrifugation at ~7000g for 10 minutes at room temperature, and re-suspended in 0.1 M citrate buffer (0.06 M citric acid- C_6_H_8_O_7_ and 0.04 M sodium citrate dihydrate- C_6_H_9_Na_3_O_9_). We repeated this procedure twice. After the final wash, we re-suspended the pellet in 1 mL LB with 20 mM sodium citrate, incubated at 37^o^C for 1 h, and pelleted and re-suspended cells in 100μl of citrate buffer and 5 mM sodium citrate. We plated cells on LB agar containing appropriate antibiotic selection, and incubated overnight at 37^o^C.

**Assaying survival under DNA damage**

We assessed strain survival under three kinds of DNA damage: exposure to UV light (base dimerization), hydrogen peroxide (base oxidation) and ciprofloxacin (double stranded DNA breaks). To assess resistance to UV, we grew cultures to saturation and then sub-cultured them 1% by volume to obtain approximately mid-log phase cultures with O.D._600_ ~ 0.6. We subjected cultures to serial dilution and plated on LB agar to obtain between 30 to 200 colonies per plate. We immediately exposed the plates to UV-C light at an intensity of 20 J/m^2^ for 5 s in a UV crosslinker (Stratagene). We used a control unexposed plate in every set in triplicate to assess mortality from UV exposure. We incubated plates for 24h at 37^o^C, wrapped in aluminium foil to prevent exposure to light and possible repair through DNA photolyase. To assess survival after exposure to hydrogen peroxide, we treated mid-log phase cultures of WT and Mutant with 5 mM hydrogen peroxide (Sigma) as described earlier ^10^. We incubated cultures at 37^o^C with shaking and monitored viable counts by plating every hour for 3 hours. We measured resistance to ciprofloxacin (“Cip”) by plating mid-log phase cultures of WT and Mutant on LB agar with 50 ng/mL Cip. However, we obtained very few resistant colonies, potentially limiting our statistical power. Previous reports indicate that prior exposure to a sub-inhibitory concentration of Cip enhances survival under a high concentration ^11^. Hence, we grew WT and Mutant to mid log phase (when they had comparable viable counts), and “pulsed” cultures with a sub-inhibitory concentration of Cip (20 ng/mL) for 1h. We used 100 µL of this culture for serial dilution and plating to measure viable counts. We centrifuged the remaining 1.9 mL culture, re-suspended the cell pellet in 100 µL LB, and plated on LB agar plates with 50 ng/mL Cip (Cip 50; inhibitory). We counted the number of colonies that arose on Cip plates after 24 hours and normalised with the total viable count taken at the same time. The difference between WT and Mutant survival was consistent whether we measured resistance by plating cultures directly on Cip 50, or after pulsing with Cip20; but the latter had higher survival overall, improving statistical power. Hence, we report data obtained from the latter set of experiments.

**Dual luciferase assay to assess miscoding**

To assess the frequency of translational errors, we used a modified version of the previously described dual luciferase assay ^12^. Briefly, the assay uses plasmids carrying fused Renilla and firefly luciferase genes encoded in the same open reading frame. Because they have distinct substrates, their activity can be independently assayed. We used the plasmid pEK7, which carries AAT (asparagine) instead of AAA (lysine) at codon 529 of the firefly luciferase gene, rendering the enzyme inactive. The assay measures how often AAT is misread as AAA, leading to incorporation of lysine at the active site and therefore a functional enzyme. Note that this method only detects decoding errors, i.e. incorrect codon-anticodon pairing. We grew cells overnight at 37^o^C in 2 mL LB medium supplemented with ampicillin (100 µg/mL) to maintain selection for the plasmid. We sub-cultured cells 1% by volume and allowed them to grow until they reached an optical density (O.D._600_) of ~0.6. We pelleted and resuspended cells in 100 µl lysis buffer (1 mg/mL lysozyme, 10 mM Tris-HCl at pH 8.0, 1 mM EDTA). After a 10-minute incubation on ice, we flash-froze cells in liquid nitrogen. We thawed them and used the whole extract for assaying firefly luciferase (F-Luc; measures miscoding) and Renilla luciferase activity (R-Luc; serves as a control) using the Dual-Luciferase Reporter assay system (Promega). For each reaction, we noted luminescence (Relative Light Units) after 10 seconds using a Wallac 1420 Victor-3 luminometer (Perkin Elmer). We calculated the final activity as the ratio of F-Luc to R-Luc activity.

**Measuring population growth rates and doubling times**

To measure growth rate across strains, we used 40 independent colonies of each *E. coli* strain as biological replicates. We inoculated colonies in LB broth and allowed them to grow overnight at 37^o^C with shaking at 200 rpm for 16 h. We added 5 µL of the overnight culture into 495 µL of the relevant (Liconic) at 37ºC. We measured OD_600_ using an automated growth measurement system (microplate reader from Tecan, Austria), every 30 or 40 minutes for 12 to 18 hours. We estimated maximum growth rates and doubling times using the Curve Fitter software (Delaney et al, 2013).

**Measuring mutation frequency using rifampicin resistance**

We estimated mutation frequency in WT and mutant cells using spontaneous mutations conferring rifampicin resistance. We grew WT and mutant to saturation (when they had comparable viable counts) at 37^o^C in a shaker incubator (Thermo Scientific). We used 100 µL of this culture for serial dilution and plating to measure viable counts. We pelleted the remaining 1.9 mL, resuspended in 100 µL LB and plated onto LB agar carrying 50 µg/mL rifampicin (Sigma Aldrich). We incubated plates for 24 h at 37^o^C and counted the number of resistant colonies. These were then divided by the total viable count to obtain the mutation frequency, as described before ^13^.

**Sanger sequencing to identify *gyrA* mutations**

To check whether the WT and mutant acquired *gyrA* mutations within 2 h of exposure to Cip, we monitored WT and mutant cultures (n=2) after exposure to Cip50 for 2 h. We pelleted cells from the whole culture after 2 h and extracted genomic DNA. We PCR amplified the ~2.5 kB *gyrA* locus using the following primers: fp 5’ GACAAACGAGTATATCAGGCAT 3’ and rp 5’ GCCACATTCCTTGTGTATAGCCACC 3’. We purified the PCR product using a PCR purification kit (Promega) followed by Sanger sequencing of the region containing the QRDR locus, using the primer 5’ GTACTACCTGACCGAACAGCAAGC 3’.

**Measuring protein production**

To measure levels of candidate proteins responsible for the observed cellular response to stress, we used western blots with a chemiluminescent detection system. We prepared cell extracts from mid log phase cultures (O.D_600nm_ ~0.6). We pelleted cells and resuspended them in sonication buffer (50 mM Tris-Cl and 150 mM NaCl with 1:100 Protease inhibitor from Genetix). We then carried out sonication (2 s on, 2.5 s off) while the samples remained in ice, with a 38% duty cycle using a micro probe sonicator . We spun down cell debris at ~10,000 g for 20 min and used the supernatant as the sample. We estimated protein concentration in each sample using Qubit (Invitrogen). We loaded 15 to 20 µg total protein on a 12% SDS polyacrylamide and ran the gel for 1.5 to 3 h (1.5 for LexA and RecA and 3 h for Lon blots) at 130V (Biorad). We then transferred the gel onto a PVDF membrane (Amersham Hybond) using electroblotting (Biorad) with Tris Glycine buffer carrying 10% methanol. For Western blotting, we used antibodies as specified in the main Methods. We used 5% non-fat milk (Genetix) overnight at 4^o^C for blocking, and treated blots with antibody for 1 h at room temperature. We used three washes after primary and secondary antibody treatment, each 15 minutes in TBST (20 mM Tris-Cl and 150 mM NaCl with 1% Tween 20). We developed blots using a chemiluminescence detection kit (Thermo Pierce) and ImageQuant LAS4000 (GE). We normalized each band of interest to the total protein (calculated using Coomassie blue stained bands of a different size from the same gel). While this allowed us to precisely account for loading differences in total protein, it introduced large variability across replicates due to differences between the sensitivity of chemiluminescence (blot) and Coomassie staining (gel).

**Assaying persister cells after exposure to antibiotics**

In addition to antibiotic resistance (a measure of active survival), we also quantified persistence under high antibiotic concentrations. We first determined that 200 ng/mL Cip served as a lethal concentration (no Cip-resistant colonies were observed on agar plates, either for WT or the Mutant). However, a few persister cells could potentially survive a brief lethal Cip concentration and subsequently form colonies in the absence of the antibiotic. To estimate the number of persister cells, we grew 2 mL cultures of WT and Mutant to mid-log phase and treated them with 200 ng/mL Cip for 2 h. We then pelleted, re-suspended and plated cells on LB agar plates without any antibiotic. We counted the number of colonies after 24 h as the number of persister cells that survived the lethal Cip treatment.

**Whole genome sequencing**

To identify mutations responsible for ciprofloxacin resistance, we inoculated 5mL LB with colonies obtained on Cip50 plates, and allowed them to grow for ~6 h (OD_600_~0.4). We extracted genomic DNA (GenElute Bacterial Genomic DNA kit, Sigma-Aldrich) and quantified it (Qubit HS dsDNA assay kit, Invitrogen). We pooled equal amounts of genomic DNA from 2 colonies per strain. For persisters obtained for 6 different strains (WT, Mutant, WT hyper-accurate, Mutant hyper-accurate, WT (no SOS) and Mutant (no SOS) colonies), we pooled equal amounts of genomic DNA from 10 persister colonies per strain. We prepared libraries (Illumina Nextera Library Preparation Kit) and added unique adaptors to identify each sample. We sequenced libraries on the Illumina Mi-seq platform (2x250 and 2x300 paired-end V2 reaction chemistry). We used the Breseq pipeline ^14^ with default parameters to identify mutations. For persisters, we obtained ~30X coverage per strain; for resistant strains, we obtained ~55–80X coverage per strain. Because the whole genome sequence of KL16 is not available, we aligned reads to the NCBI reference *E. coli* K-12 MG1655 genome (RefSeq accession ID GCA_000005845.2). We also sequenced the WT KL16 genome and compared it against the MG1655 genome to identify and filter out KL16-specific mutations. We only called mutations that were supported by at least 20 reads (5 reads for pooled persister samples) and with >40% frequency.

**SUPPLEMENTARY FIGURES AND TABLES**

**Figure S1:** **Phenotype microarray for WT and Mutant**. WT and Mutant were inoculated in the phenotype microarray antibiotic plate PM12B. Each antibiotic is present in 4 wells, with 1X, 2X, 4X and 8X dilutions in successive wells. Absolute concentrations are unknown. The heat map shows the difference in the area under the curve (AUC) of growth of Mutant minus WT after 48h. The three antibiotics with the highest differences are highlighted.

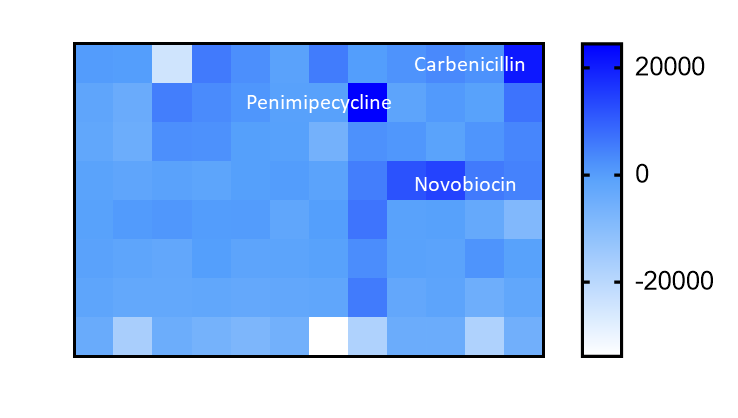

**Figure S2:** **(a) Mutant grows slower than the WT**. Mean doubling time of WT and Mutant cultures (n=44) estimated from the growth curve. Mann-Whitney U test: WT >Mutant, U=4, P<0.0001 **(b)** **Reducing WT growth rate does not impact ciprofloxacin resistance**. Mean survival of WT and Mutant cultures from single colonies (n=6) pulsed with 20 ng/mL ciprofloxacin (Cip 20) for 1 h and plated on LB agar with vs. without 50 ng/mL Cip (Cip 50). Plot shows the number of resistant colonies per unit viable count from LB plates. Mann-Whitney U test: WT vs WT (glycerol), ns, U=1, P=0.07. Asterisks indicate significant differences.

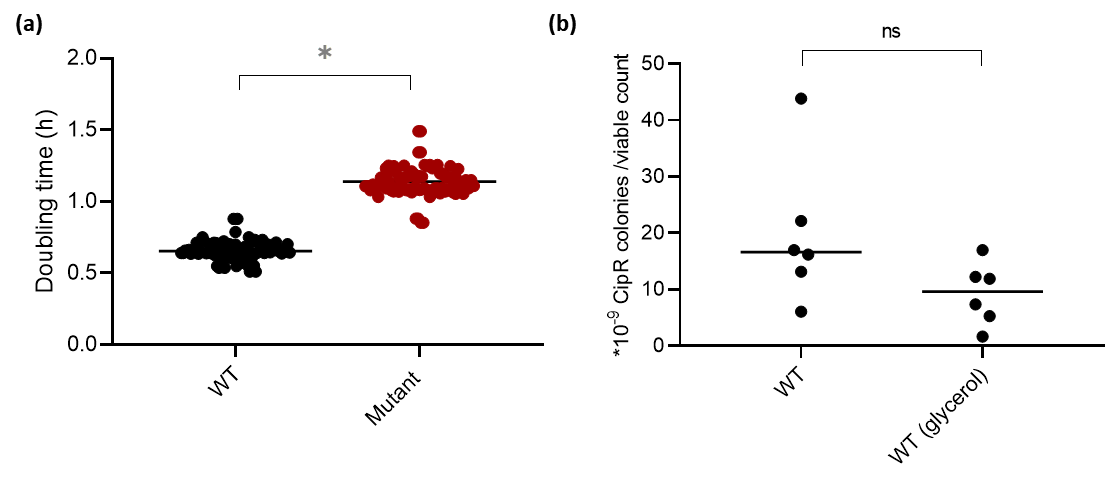

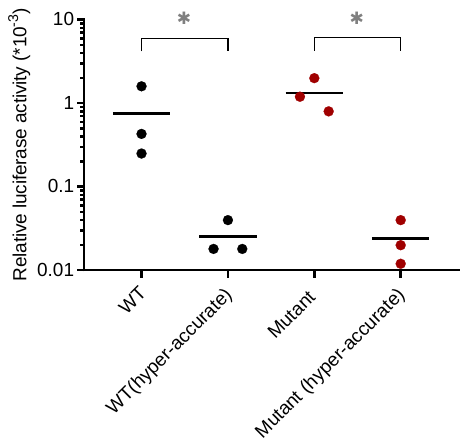

**Figure S3**: **WT and Mutant show similar mutation frequencies.** Overnight cultures from single colonies of WT and Mutant were plated on LB agar with 50 µg/mL of rifampicin (n=15, with paired control plates for viable counts). Plot shows the mean number of resistant colonies per unit viable count from LB plates. WT vs Mutant, ns, Mann-Whitney U test, U=100.5, P=0.63.

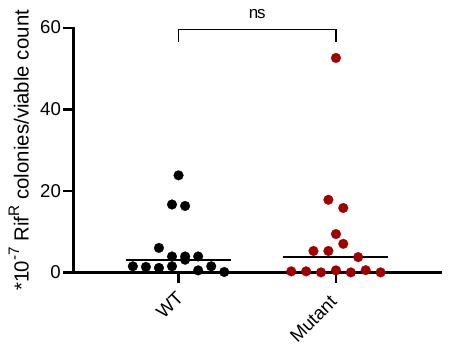

**Figure S4**: **Hyper-accurate strains show lower mistranslation.** Mean mistranslation rates measured with an *in vitro* dual luciferase assay for WT and mutant (n=3). Asterisks indicate significant differences. Paired t tests: WT (hyper-accurate)<WT, t=6.9, P=0.006; Mutant (hyper-accurate)<Mutant ,t=4.8, P=0.008.

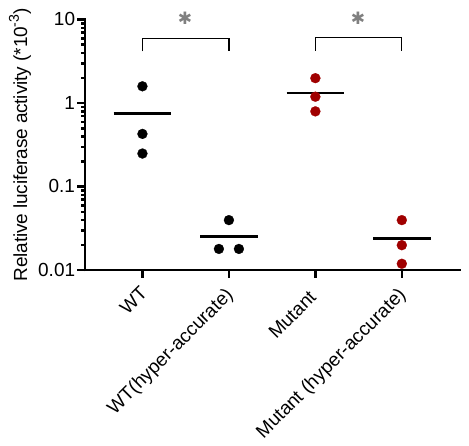
.

**Figure S5**: **Inactivating the SOS response reduces early survival in ciprofloxacin.** Mean survival of WT and Mutant cultures from single colonies (n=3) treated with Cip 50 for 2 h and plated on LB agar. Plot shows the percent survival calculated from total viable counts before and after Cip treatment. Asterisks indicate significant differences. SOS was inactivated using the *lexA3* allele. t tests: WT(inactive SOS)<WT, t=6.2, P=0.02; Mutant(inactive SOS)<Mutant, t=7.8, P=0.02; WT (inactive SOS) vs. Mutant (inactive SOS), ns, t=0.96, P=0.43.

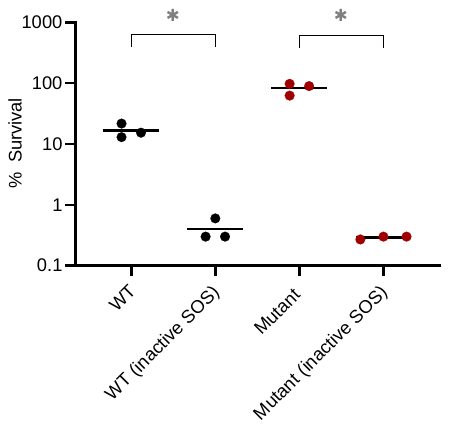

**Figure S6: Deletion of RecN reduces early survival in ciprofloxacin.** Survival of mid log phase cultures (OD_600nm_~0.6) of the indicated strains from single colonies (n=3) treated with Cip50 for 2 h and plated on LB agar. Plot shows the mean % survival calculated from total viable counts before and after Cip treatment. t test: WΔ*recN* <WT, t=4.9, P=0.03; MutantΔ*recN*<Mutant, t=7.6, P=0.02; WTΔ*recN* vs MutantΔ*recN* , ns, t=0.96, P=0.43.

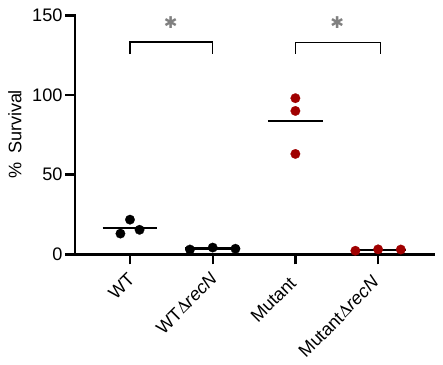

**Figure S7: Time course of LexA degradation by WT and mutant.** LexA protein levels normalised to total protein as measured by western blotting using a polyclonal anti-LexA antibody. Each panel shows data for an independent experimental block, in addition to the block shown in Fig.2e.

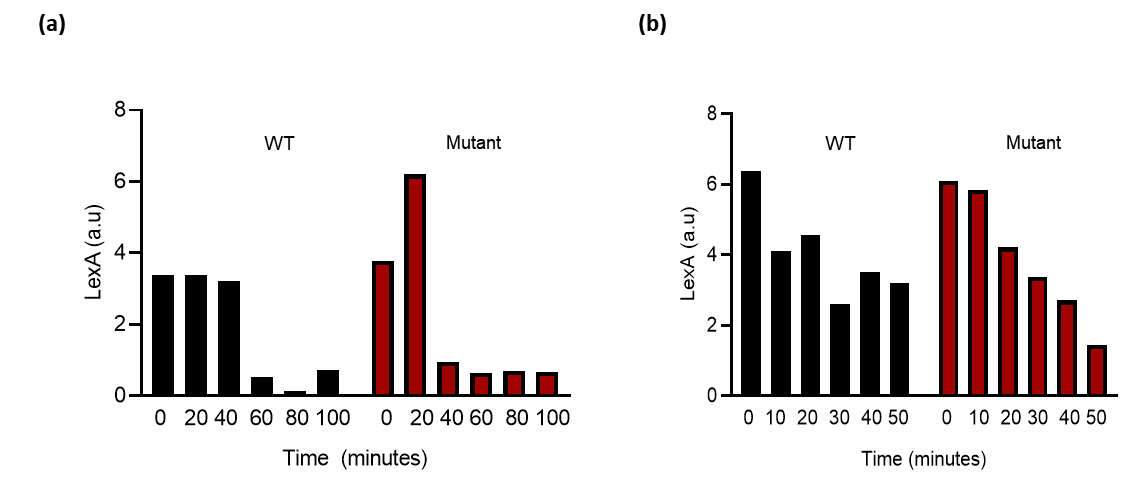

**Figure S8**: **The mutant activates SOS at lower levels of stress.** (a) LexA protein levels in WT and mutant cultures exposed to different Cip concentrations for 30 mins, measured by western blotting using a polyclonal anti-LexA antibody. Quantitation across biological replicates (n=3) is shown (mean±SD). (b) RecA protein levels in WT and mutant cultures, measured by western blotting using a polyclonal anti-RecA antibody. Quantitation across biological replicates (n=3) is shown (mean±SD).

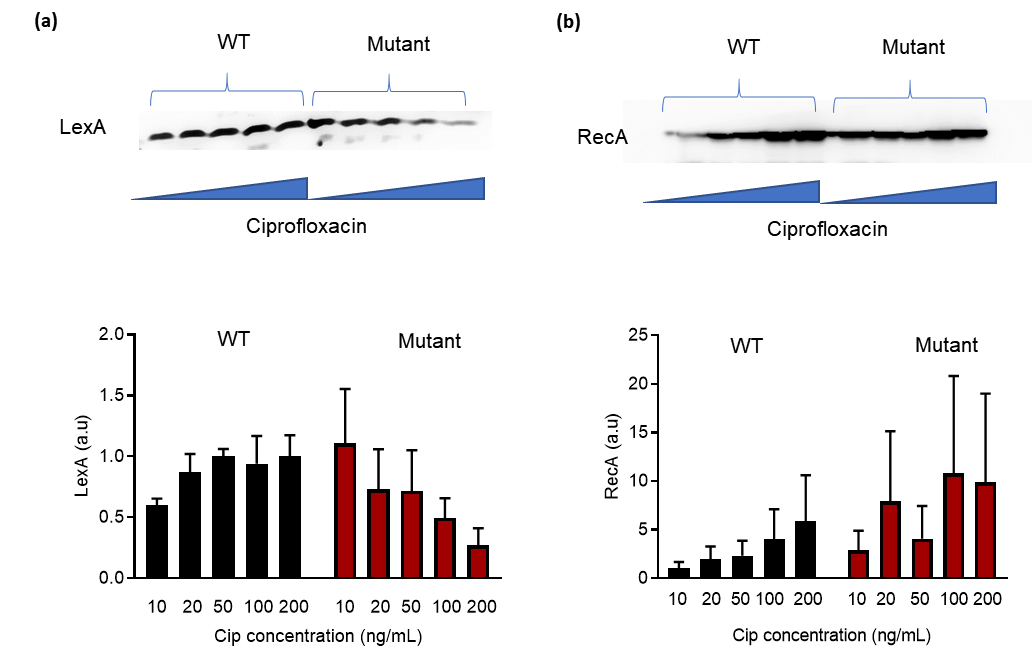

**Figure S9: Mistranslation elevates the heat shock transcription factor sigma 32.** Mean levels of Sigma 32 protein in mid log phase cultures (OD_600_~0.6) assessed by western blotting using a polyclonal anti-sigma 32 antibody. Quantitation across biological replicates (n=3) is shown. Paired t tests: Mutant vs. WT, ns, t=3.1, P=0.08; WT (canavanine) vs. WT, ns, t=4.08, P=0.05; WT(norleucine)>WT, t=4.5, P=0.04.

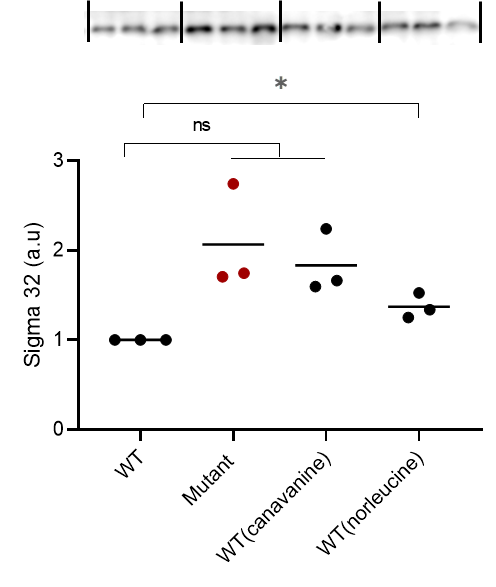

**Figure S10**: **WT (KL16) and MG1655 show comparable ciprofloxacin resistance.** Resistance of mid log phase cultures of WT (KL16) and MG1655 (OD_600nm_~0.6) from single colonies (n=6) pulsed with Cip 20 for 1 h and plated on LB agar with vs. without Cip 50. Plot shows the mean number of resistant colonies per unit viable count from LB agar plates. Mann-Whitney U test, WT vs MG1655, ns, U=10, P=0.76.

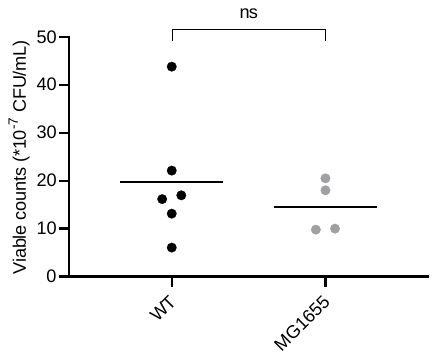

**Figure S11: Over-expression of Lon protease increases early survival in ciprofloxacin.** Early survival (tolerance) of mid log phase cultures (OD_600nm_~0.6) of WT and WT+Lon from single colonies (n=3) treated with Cip 50 for 2 h and plated on LB agar. Plot shows the mean % survival calculated from total viable counts before and after Cip treatment. Paired t test: WT+Lon>WT, t=5.1, P=0.03

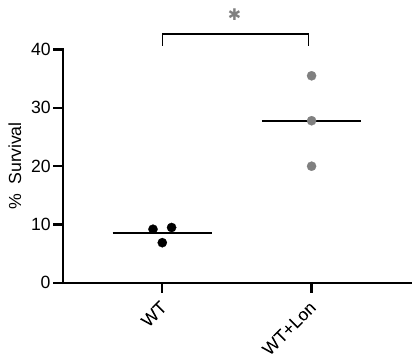

**Figure S12: Over-expression of Lon protease elevates RecA levels.** RecA protein levels in mid log phase (OD_600_~0.6) cultures assessed by western blotting using a polyclonal anti-RecA antibody, normalised to total protein. The figure shows bands from two separate blots.t test: WT+Lon>WT, t=5.13, P=0.03.

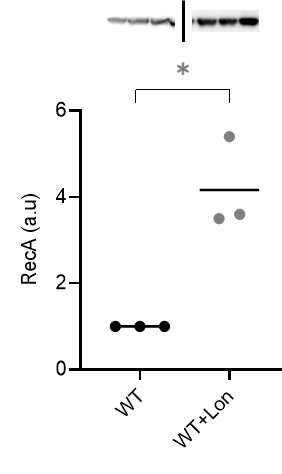

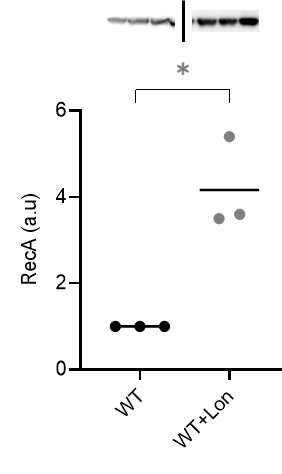
**`**

**Figure S13: Mistranslation provides an early but variable survival advantage under high temperature.** Survival of cultures inoculated from single colonies (n=3), dilution plated at indicated times on LB agar. Two experimental blocks are shown. Plot shows the total viable counts (mean±SEM). (a) t-tests at 24 h: Mutant>WT, t=7.2, P<0.001; WT(norleucine)>WT, t=5.6, P= 0.007; WT vs WT(canavanine), ns, t=0.12, P=0.9 (b) t-tests at 24 h: WT vs Mutant, ns, t=2.8, P=0.06; WT(canavanine)>WT, t=3.1, P=0.02.

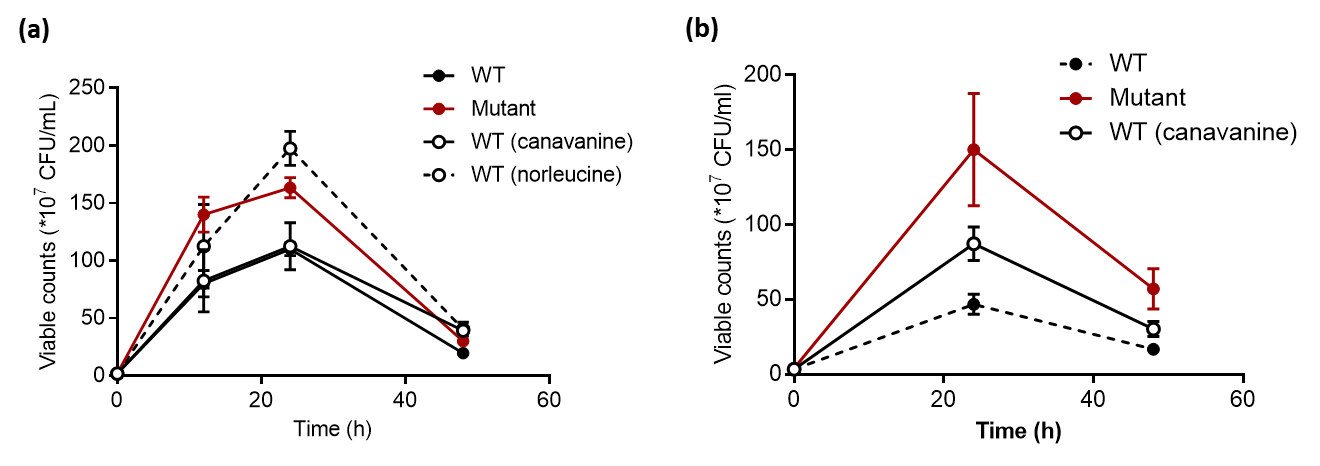

**Figure S14:** **WT and mutant have comparable Lon levels under heat stress.** Mean levels of Lon protein in mid log phase (OD_600_~0.6) cultures of WT and Mutant (n=3) as measured by western blotting using a polyclonal anti-Lon antibody, normalised to total protein. Paired t test: WT vs Mutant, ns, t=0.9, P=0.12.

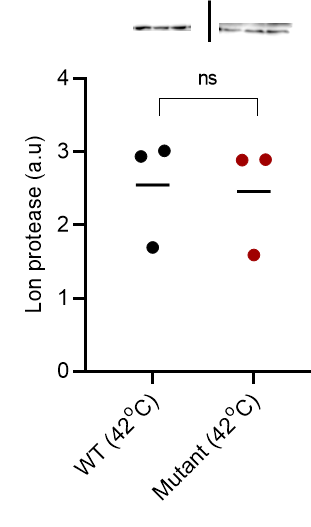

**Figure S15:** **Mutant degrades LexA under heat stress.** Mean LexA protein levels in mid log phase (OD_600_~0.6) cultures of WT and Mutant (n=3) as measured by western blotting using a polyclonal anti-Lon antibody, normalised to total protein. Paired t test: Mutant<WT, t=5.3, P=0.03.

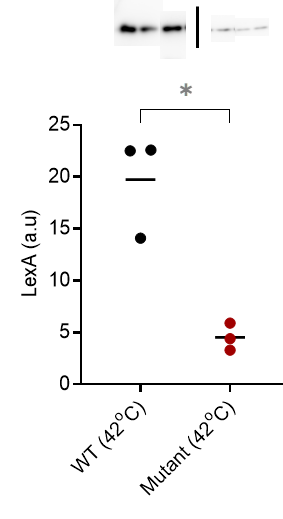

**Figure S16:** **RecN provides an early survival advantage under high temperature.** Survival of cultures inoculated from single colonies (n=3 for WTΔr*ecN* and MutantΔr*ecN* and n=5 for WT and Mutant), dilution plated at indicated times on LB agar. Plot shows the total viable counts (means±SEM). Unpaired t-tests at 12 h: WTΔr*ecN*<WT, t=17.8, P<0.0001; Mutant*ΔrecN*<Mutant, t=5.2, P= 0.002. At 24 h: WT*ΔrecN* vs. WT, ns t=0.79, P=0.28; Mutant*ΔrecN <*Mutant, t=5.2, P= 0.002.

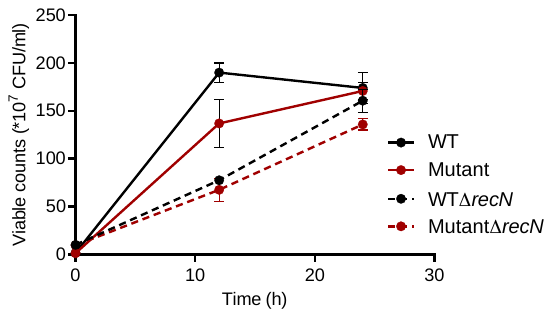

**Table S1: Mutations in ciprofloxacin-resistant WT and mutant colonies after 24 h on a Cip 50 plate.** Numbers in parentheses indicate the number of individual colonies carrying each mutation.

| **WT (5)** | **Mutant (5)** |
| --- | --- |
| *gyrA*-S83L (2) | *gyrA*-S83L (3) |
| *gyrA*-G81D (2) | *gyrA*-D87G (2) |
| *gyrA*- D87G (1) |  |
